# Duplication and losses of opsin genes in lophotrochozoan evolution

**DOI:** 10.1101/2022.07.29.501998

**Authors:** Giacinto De Vivo, Fabio Crocetta, Miriam Ferretti, Roberto Feuda, Salvatore D’Aniello

## Abstract

Opsins are G-coupled receptors playing a key role in metazoan visual processes. While many studies enriched our understanding of opsin diversity in several animal clades, the opsin evolution in Lophotrochozoa, one of the major metazoan groups, remains poorly understood. We investigated the opsin evolution in 74 translated lophotrochozoan genomes and capitalized on recently developed phylogenetic approaches. We found that the common ancestor of Lophotrochozoa possessed at least seven opsin paralog groups that underwent divergent evolutionary history in the different phyla. Furthermore, we showed for the first time placopsin-related molecules in Bilateria, that we named pseudopsins, which may prove critical in uncovering opsin evolution.

## Introduction

Lophotrochozoa is one of the largest metazoan clades and includes annelids, molluscs, nemerteans, bryozoans, brachiopods, and platyhelminths (Luo et al. 2018; Marlétaz et al. 2019). Species of this group occupy many ecological niches in aquatic and terrestrial environments and exhibit diverse photoreceptive visual structures, from simple pigmented areas to sophisticated camera-type eyes (Serb and Eernisse 2008; Bok et al. 2017; Rawlinson et al. 2019). Yet, the evolution and distribution of the opsin genes in this clade remain poorly understood.

Opsins mediate the initial step in visual processes in almost all animals. Using a highly conserved lysine (K296 with reference to the bovine rhodopsin), they bind using a Schiff-linkage to the retinal, a vitamin A derivative, which acts as a chromophore. When the retinal absorbs a photon, it isomerises from 11-*cis* to all-*trans*, causing conformational changes to the opsin, which activate the G protein to initiate the signalling cascade (Terakita 2005; Shichida and Matsuyama 2009). Amino acid sequence variation surrounding the retinal molecule can shift the optimal wavelength absorption (Shichida and Imai 1998). The increase in the opsin complement is associated with better visual capabilities, although this view has been recently challenged, and new physiological roles in mechanoreception, chemoreception, thermoception, and cellular membrane regulation have been proposed (Feuda et al. 2022).

Opsins originated early in metazoans, and three major lines, ciliary (c-opsin), Group-4 (RGR/Go) opsins, and rhabdomeric (r-opsin), were already present in the Cnidaria-Bilateria ancestor (Feuda et al. 2012; Picciani et al. 2018; Fleming et al. 2020). This complement further expanded in Bilateria, where nine subfamilies were described: r-opsins, non-canonical r-opsins, c-opsins, xenopsins, chaopsins, bathyopsins, retinochrome/RGR/peropsins, neuropsins, and Go-opsins (D’Aniello et al. 2015; Ramirez et al. 2016; Vöcking et al. 2017). The eumetazoan opsins share a common ancestor with placopsins, opsin-like molecules, which have been identified in placozoans. Differently from conventional opsins, these proteins lack retinal binding residues and might not have a photoreceptive function (Feuda et al. 2012).

Previous works have investigated the opsin evolution in lophotrochozoans focusing mostly on individual species or specific clades (Passamaneck et al. 2011; Gühmann et al. 2015; Yoshida et al. 2015; Ramirez et al. 2016; Bok et al. 2017; Vöcking et al. 2017; Rawlinson et al. 2019; Bonadè et al. 2020; Döring et al. 2020); as consequence, the complete view of the evolution and the long-term dynamics of opsin genes in this clade remains poorly understood. Here, we study the evolution of opsin genes in 11 lophotrochozoan phyla using a phylogenomics approach. Our results suggest a complex pattern of duplications and losses in the different lophotrochozoan clades.

## Results

### The phylogeny of opsin in lophotrochozoans

To investigate the opsin evolution in lophotrochozoans, we used 74 translated genomes from 11 phyla. Opsins were identified using BLASTp and organized into different ortho-groups using Broccoli (Derelle et al. 2020). The phylogenetic relationships were reconstructed using Ultrafast Bootstrap (UFB) and Transfer Bootstrap Expectation (TBE), methods specifically designed for single-gene phylogeny (Lemoine et al. 2018; Minh et al. 2020). After removing sequences with less than three transmembrane domains (see methods for more details), we performed a first phylogenetic analysis using UFB (UFB1) and then TBE (Figure 1A, Suppl. File S1 and Suppl. File S2) to identify unstable sequences (rogue taxa) that were subsequently removed from our dataset (380 putative opsin sequences retained), and used for a new UFB phylogenetic analysis (UFB2) (Figure 1B, Suppl. File S3). All phylogenetic trees support the monophyly of Go-opsins, neuropsins, peropsins, and RGR opsins, hereafter Group-4 (Figure 1) (UFB1=95; TBE=0.76; UFB2=100), as well as the sister relationship between c-opsins and xenopsins (UFB1=66; TBE=0.68; UFB2=91). C-opsins/xenopsins and Group-4 form a monophyletic group (UFB1=84; TBE=0.86; UFB2=98), which is the sister group to the r-opsins (UFB1=82; TBE=0.76; UFB2=79).

**Figure 1.**
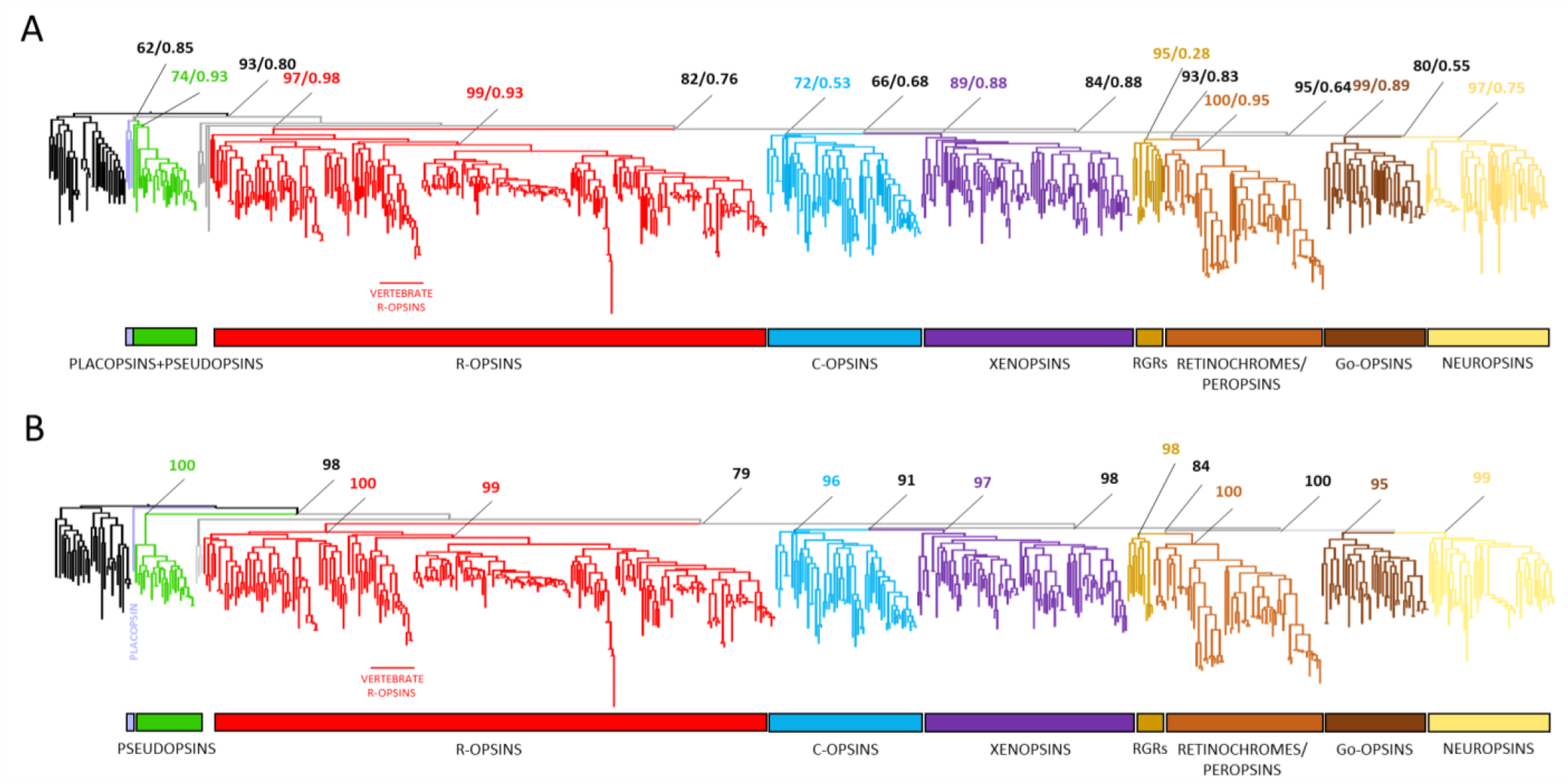
– Metazoan opsin phylogeny. The UFB1 and TBE tree is shown in A, and the UFB2 tree in B, computed after removing unstable sequences. Seven opsin groups are found in lophotrochozoans, with r-opsins being the sister group to all the other opsins, while xenopsins are strongly related to c-opsins. The opsin-related sequences, pseudopsins, clade with placopsins (A) or opsins (B). The outgroup, in black, represents melatonin receptors (MLT).

Here we identified several sequences in nemerteans, annelids, and molluscs, that are placed as the sister group to opsins (UFB2=98; Figure 1B), that we named pseudopsins for their ambiguous nature. In our TBE and UFB1 analysis, before removing rogue taxa, these sequences form a monophyletic group with placopsins (UFB1=62, TBE=0.85; Figure 1A). Similarly to placopsins, pseudopsins lack the canonical retinal binding domain, although show the presence of conserved motifs indicating G-protein binding capability (Suppl. Figure S1). Furthermore, they were assigned to a single ortho-group in the Broccoli analysis and, like placopsins, connect first to opsins in the CLANS analysis (Frickey and Lupas 2004) (Suppl. Figure S2). Additionally, opsins with a diverging retinal binding domain are not restricted to the outgroup, as we also found 12 sequences nested in the opsin tree lacking the canonical retinal binding domain, representing probably sub-functionalised opsins or sequencing errors.

### The dynamics of opsin genes in Lophotrochozoa

Next, to investigate the dynamics of opsin genes in Lophotrochozoa, we initially mapped opsin paralogs in the species analysed. After removing unstable sequences, we reconstructed the pattern of gene duplication and the ancestral node content using GeneRax, a maximum likelihood gene-tree to species-tree reconciliation method (Morel et al. 2020). Overall, our results suggest that the evolution of opsin is largely dynamic in lophotrochozoans, with several events of lineage-specific duplications and losses (Figure 2A-B and Figure 3).

**Figure 2.**
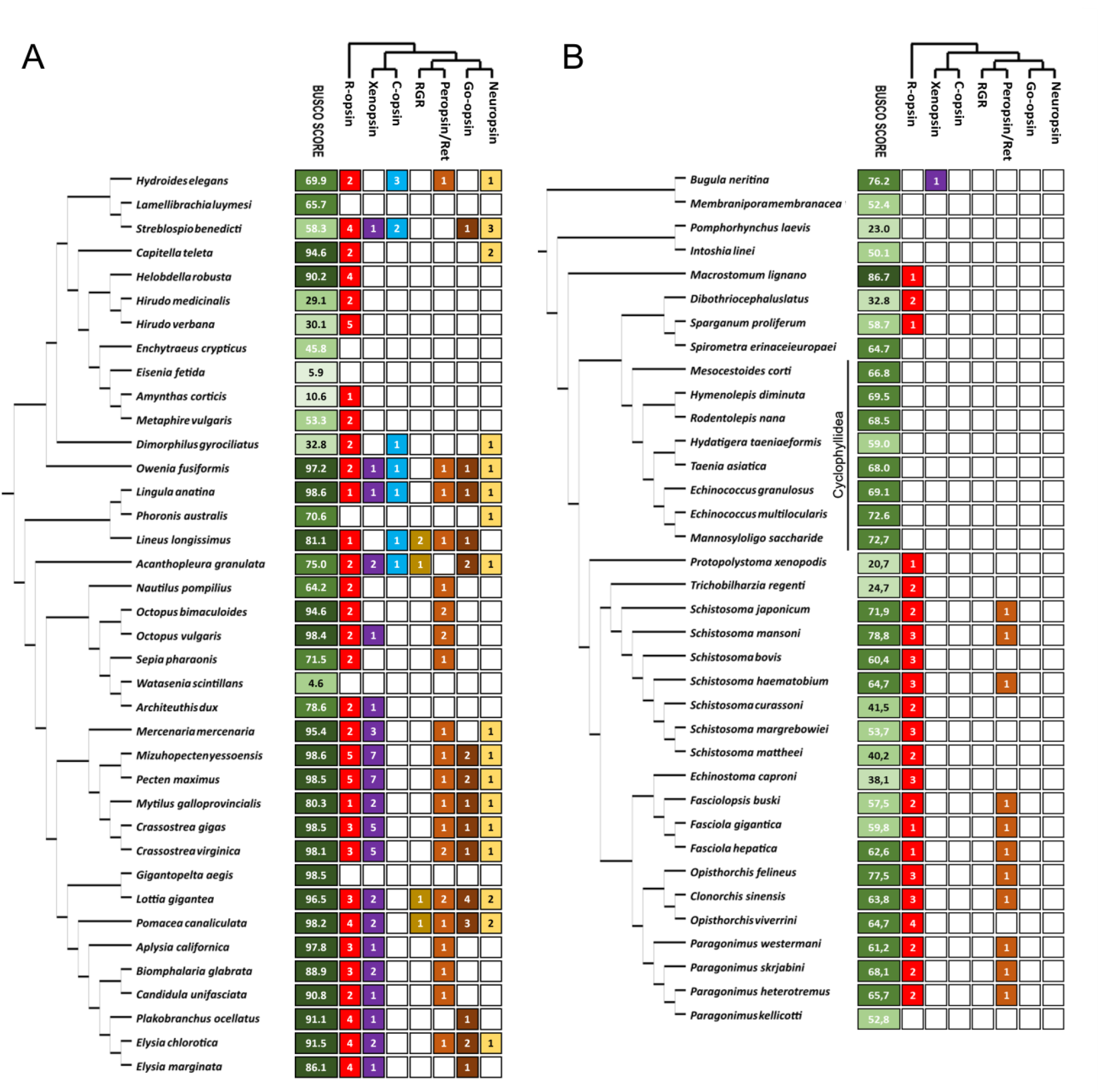
Species tree and gene tree reconciliation. The computed species tree, the minimal number of opsin found per species (species-specific duplications are not included), and the estimated number of opsins in the major clades are shown in the two cladograms, showing respectively the Annelida-Mollusca node (A) and the Bryozoa-Platyhelminthes node (B). Species named as deposited in NCBI.

**Figure 3.**
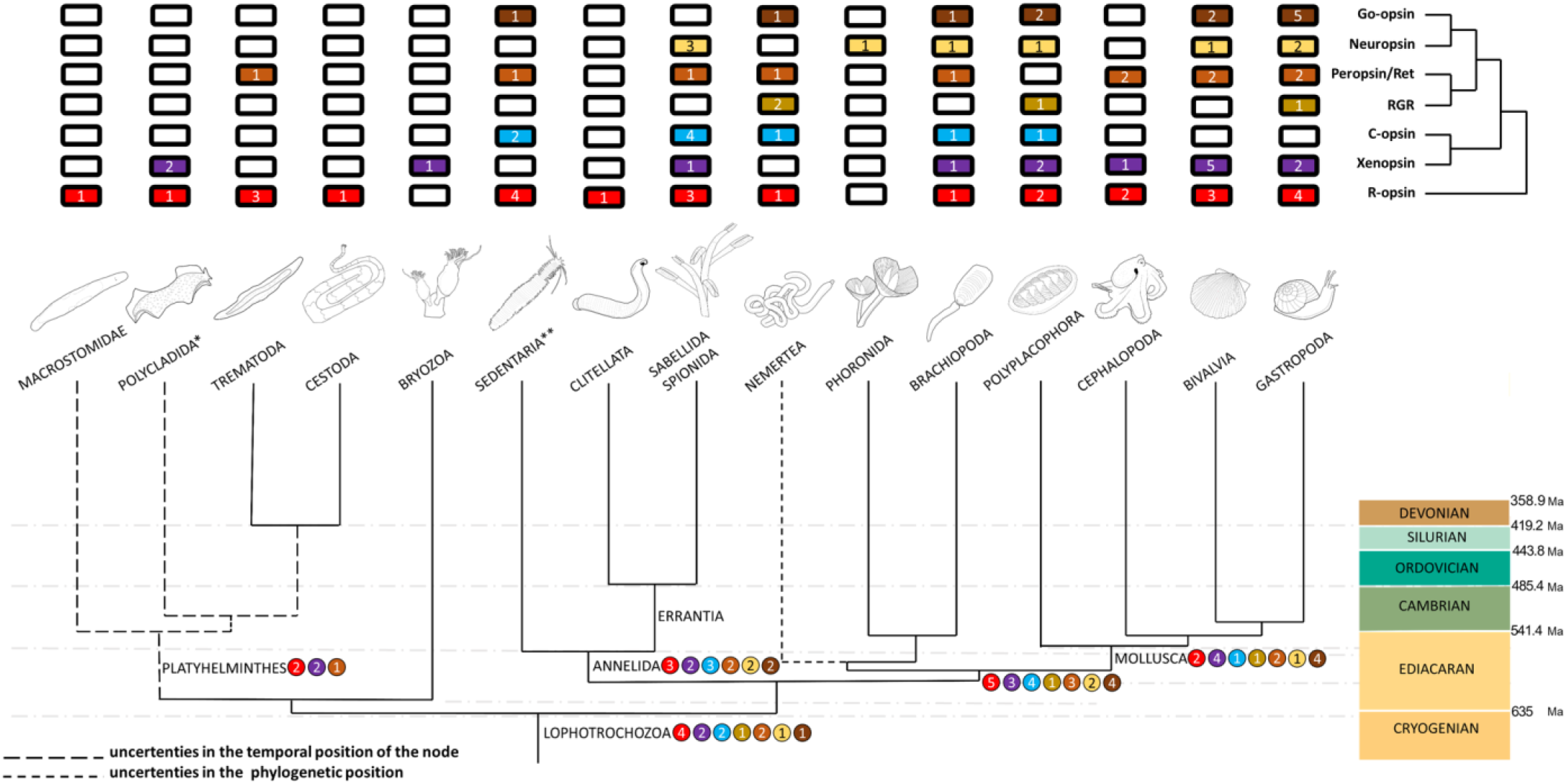
– Pattern of opsin duplications and losses in Lophotrochozoa. The estimated number of opsins per represented taxa is shown, including the reconstruction of the number of opsins at the nodes. The common ancestor of lophotrochozoans possessed a large set of opsins compared to modern representatives. A pattern of radical opsin loss can be found in the lineages leading to platyhelminths, bryozoans, clitellates, and phoronids. The symbol * indicates data from Rawlison et al. (2019) and ** data from Gühmann et al. (2015). Animal silhouettes are made from Wikipedia Commons photos, credits go to Dita Vizoso, Betty Wills (Atsme), Martin Gühmann, Karl Ragnar Gjertsen, OAR/National Undersea Research Program, Kirt L. Onthank, Hectonichus, Stako, and own work.

The opsin complement estimated from GeneRax suggests that at least 13 opsins, from seven paralog groups, were already present in the lophotrochozoan common ancestor: four r-opsins, two c-opsins, retinochrome/peropsins, and xenopsins, one RGR, Go-opsin, and neuropsin (Figure 3). Furthermore, our results suggest that, starting from the ancestral opsin complement, the bryozoan and platyhelminth lineages underwent a radical reduction of opsin genes (Figure 2 and Figure 3). Modern bryozoans only retained xenopsins, while all platyhelminths only possess r-opsins and retinochromes, although it has been shown the expression of a xenopsin in larval eyes and ciliary structures (phaosomes) of polycladids, indicating that they were present in the platyhelminths common ancestor (Rawlinson et al. 2019). Interestingly, among the analysed cestodes, a group of parasitic worms, only pseudophyllideans possess opsins, while cyclophyllideans, prevalently parasites of humans and domesticated animals, do not (Kikuchi et al. 2021).

Annelids, molluscs, nemerteans and brachiopods did not undergo a dramatic reduction of the opsins genes, in contrast to bryozoans and platyhelminths. Indeed, they retain a similar complement to the lophotrochozoan ancestor followed by specific gene duplications and independent opsin losses in many clades (Figure 3). Remarkably, we highlight that annelids lack RGRs opsins, and in particular clitellates have drastically reduced their opsin content, preserving only r-opsins. The presence of one Go-opsin, four r-opsins, and two c-opsins has been previously reported in *Platynereis dumerilii* (Gühmann et al. 2015). Brachiopods lost RGRs, and nemerteans show the presence of RGRs, retinochromes, Go-opsins, c-opsins, and r-opsins. Phoronids only possess a neuropsin and a putative RGR, the last recognised as an unstable sequence by the TBE analysis, and subsequently discarded (Figure 3). The molluscan common ancestor retained the entire opsin complement; nevertheless, only polyplacophorans (chitons) possess a c-opsin. Furthermore, bivalves and gastropods show multiple xenopsin paralogues. Surprisingly, cephalopods present the lowest number of opsins among molluscs, retaining only two r-opsins, one xenopsin, and two peropsin/retinochromes (Figure 2), and this is in contrast with the expectation given the morphological complexity of their eyes.

## Discussion

Our results substantially clarify the evolution of the opsin genes in lophotrochozoans over 600 million years, suggesting a diverging evolutionary history of opsins in this clade (Figure 3). Overall, our phylogenetic tree supports the sister relationship of c-opsins and Group-4 opsins, with r-opsins being sister to this clade, similarly to previous works (Yoshida et al. 2015; Vöcking et al. 2017; Rawlinson et al. 2019; Bonadè et al. 2020; Fleming et al. 2020). Noteworthy, our tree supports the monophyly of xenopsins and c-opsins similarly to some of the studies (Yoshida et al. 2015; Vöcking et al. 2017) but differently from others (Ramirez et al. 2016; Rawlinson et al. 2019; Bonadè et al. 2020). Remarkably, we identified in lophotrochozoans a novel bilaterian opsin-like clade related to placopsins, that we here named pseudopsins. Pseudopsins lack the retinal binding domain, and therefore possibly the photoreceptive function; however, the presence of G-binding conserved domains suggests that they might be functional. The presence in bilaterians of sequences related to placopsins would indicate a large distribution of these genes in animals, suggesting the need for a re-evaluation of opsin evolutionary history and encouraging new studies investigating the expression pattern and function of these cryptic receptors in the future.

Moreover, our results suggest the presence of a rich opsin repertoire composed of seven paralogs in the last common lophotrochozoans ancestor, in agreement with Ramirez et al. (2016). The observation of the opsin complement in the species tree (Figure 3) suggests two opposite tendencies: (i) the opsin reduction in the platyhelminths and bryozoans, and (ii) a stable number of opsins in the ancestors of molluscs, brachiopods, nemerteans, and annelids. This indeed raises the question on which are the evolutionary forces that shaped opsin diversity in lophotrochozoans. The adaptation to the parasitic lifestyle in some platyhelminth lineages might partially explain the reduced opsin complement in this clade. In other cases, environmental factors, such as the terrestrialization in clitellates, or life in extreme habitats, such as the hydrothermal vents in the deep sea for *Lamellibrachia luymesi* and *Gigantopelta aegis*, may have led to the loss of opsins (Lan et al. 2021). Furthermore, events of genome compaction could represent further causes of opsin number reduction, such in the case of *Dimorphilus gyrociliatus* (Martín-Durán et al. 2021). Importantly, we observe also that the opsin complement does not correlate with the morphological complexity of the eyes. For example, cephalopods massively reduced their opsin toolkit compared to other molluscs, despite possessing complex camera-type eyes and well-known visual capabilities. Noteworthy, *Nautilus pompilius* has the lowest number of opsins among molluscs, retaining only two r-opsins and one retinochrome (Zhang et al. 2021).

The correlation of the opsin number with complex lifecycles or with the number of photoreceptive structures might explain some differences between the opsin content in the different taxa. Furthermore, it is possible that the rich repertoire of opsins was employed for biological purposes other than vision (Feuda et al. 2022). Remarkably, the opsin function in lophotrochozoan remains poorly understood and additional work is required to understand the diversity of functions, including the role of the enigmatic xenopsins (Arendt et al. 2017), largely duplicated in bivalves and gastropods.

In conclusion, based on our results, lophotrochozoans might represent excellent models for studying the evolution of photoreception in animals, as shown by their plethora of opsins, visual structures and lifecycles. However, while our work clarifies the pattern of opsin evolution in lophotrochozoans, it still suffers from the limited genomic information available for certain groups (such as bryozoans, nemerteans, brachiopods, and phoronids) and the lack of chromosomal scale assembly for most lophotrochozoan species. The improvement of genome quality will help to refine the details of opsin duplications and further work will clarify the expression pattern and the function of different opsin genes.

## Materials and Methods

### Data mining

We downloaded 15 lophotrochozoan genomes and 59 proteomes from NCBI and UniProt (TheUniProtConsortium 2020; Rangwala et al. 2021) and analysed them as described in Feuda et al. (2016), with few modifications, as indicated below. Proteomes were extracted from selected genomes using Augustus (training sequence set: *Homo sapiens*) (Stanke et al. 2004). To create the opsin-dataset, proteomes were mined using BLASTp (e-value of 1E^−10^) and the dataset from Feuda et al. (2012) was used as a query. The resultant sequences were divided into ortho-groups using Broccoli (Derelle et al. 2020), and each sequence within the ortho-group was annotated using EggNog (Huerta-Cepas et al. 2019). Ortho-groups containing annotated opsins were added to the opsin dataset. Given the low quality of their genomes, we could not assess the presence of opsins in Acanthocephalans and Orthonectids.

TOPCONS (Bernsel et al. 2009) was used to identify the number of transmembrane domains, and sequences with less than three transmembrane domains were removed. Two distinct datasets were created: (i) “opsin dataset 1” used for gene phylogeny, containing all the sequences found in our analysis plus a pre-compiled list of annotated opsins from the literature (Suppl. File S4), and (ii) the “opsin dataset 2” used for GeneRax analysis, in which the opsin outgroups (pseudopsins and melatonin receptors) and unstable sequences were removed (Suppl. File S5). In parallel, the single-copy orthologs dataset for each species was extracted with BUSCO (lineage dataset: metazoa_odb10; Seppey et al. 2019).

### Phylogeny and tree reconciliation

The final and the unrooted opsin datasets have been aligned in MEGA using the MUSCLE algorithm (Kumar et al. 2008). Single-copy orthologs were aligned using MAFFT and assembled with FasconCAT (Kück and Longo 2014; Katoh et al. 2017). We first run on IQTREE.2 an initial UFB analysis (UFB1, GTR20+G4 model with 1000 UBF replicates) and then a TBE (transfer bootstrap expectation) analysis (GTR20+G4 model 100 bootstrap replicates) using opsin dataset 1 to spot rogue taxa (unstable sequences with a transfer bootstrap index over 95% quantile, Suppl. File S6) (Lemoine et al. 2018; Minh et al. 2020). Once removed the unstable sequences from opsin dataset 1, we repeated the UFB phylogenetic analyses (UFB2). All the gene trees were rooted using melatonin receptors (Feuda et al. 2012).

To conduct the GeneRax analysis, the species tree and the gene tree have been computed using respectively the single-copy orthologs dataset and the opsin dataset 2 on IQTREE (GTR20+G4 model with 1000 UBF). The species tree was rooted using the split Annellida-Gastropoda and Bryozoa-Platyhelminthes (Kocot et al. 2016; Luo et al. 2018). We manually corrected the phylogenetic position of Dinophillidae, Oweniidae, and Nemerteans using MesquiteX.X (Maddison and Maddison 2017; Carrillo-Baltodano et al. 2021; Martín-Durán et al. 2021). The species tree and the unrooted gene tree have been subsequently reconciled using GeneRax with PROTGTR+G4 substitution model and *Danio rerio* as outgroup (Morel et al. 2020). Finally, the pattern of gene loss and duplication has been explored and annotated.

## Acknowledgements

We thank Clifton Lewis, Alessandra Aleotti, Julien Devilliers, and Matthew Goulty for their bioinformatic help and support. We also thank John Kirwan for critical reading of the manuscript and his help with the R script.

## Supplementary Material

Supplementary material is available on https://

## References

Arendt D. 2017. The enigmatic xenopsins. eLife 6:e31781.

Bernsel A, Viklund H, Hennerdal A, Elofsson A. 2009. TOPCONS: consensus prediction of membrane protein topology. Nucleic Acids Research 37:W465–W468.

Bok MJ, Porter ML, Ten Hove HA, Smith R, Nilsson D-E. 2017. Radiolar Eyes of Serpulid Worms (Annelida, Serpulidae): Structures, Function, and Phototransduction. The Biological Bulletin 233:39–57.

Bonadè M, Ogura A, Corre E, Bassaglia Y, Bonnaud-Ponticelli L. 2020. Diversity of Light Sensing Molecules and Their Expression During the Embryogenesis of the Cuttlefish (Sepia officinalis). Frontiers in Physiology 11:521989.

Carrillo-Baltodano AM, Seudre O, Guynes K, Martín-Durán JM. 2021. Early embryogenesis and organogenesis in the annelid Owenia fusiformis. EvoDevo 12:5.

D’Aniello S, Delroisse J, Valero-Gracia A, Lowe EK, Byrne M, Cannon JT, Halanych KM, Elphick MR, Mallefet J, Kaul-Strehlow S, et al. 2015. Opsin evolution in the Ambulacraria. Marine Genomics 24:177–183.

Derelle R, Philippe H, Colbourne JK. 2020. Broccoli: Combining Phylogenetic and Network Analyses for Orthology Assignment. Molecular Biology and Evolution 37:3389–3396.

Döring CC, Kumar S, Tumu SC, Kourtesis I, Hausen H. 2020. The visual pigment xenopsin is widespread in protostome eyes and impacts the view on eye evolution. eLife 9:e55193.

Feuda R, Hamilton SC, McInerney JO, Pisani D. 2012. Metazoan opsin evolution reveals a simple route to animal vision. Proceedings of the National Academy of Sciences 109:18868–18872.

Feuda R, Marlétaz F, Bentley MA, Holland PWH. 2016. Conservation, Duplication, and Divergence of Five Opsin Genes in Insect Evolution. Genome Biology and Evolution 8:579–587.

Feuda R, Menon AK, Göpfert MC. 2022. Rethinking Opsins. Molecular Biology and Evolution 39:msac033.

Fleming JF, Feuda R, Roberts NW, Pisani D. 2020. A Novel Approach to Investigate the Effect of Tree Reconstruction Artifacts in Single-Gene Analysis Clarifies Opsin Evolution in Nonbilaterian Metazoans. Genome Biology and Evolution 12:3906–3916.

Frickey T, Lupas A. 2004. CLANS: a Java application for visualizing protein families based on pairwise similarity. Bioinformatics 20:3702–3704.

Gühmann M, Jia H, Randel N, Verasztó C, Bezares-Calderón LA, Michiels Nico K, Yokoyama S, Jékely G. 2015. Spectral Tuning of Phototaxis by a Go-Opsin in the Rhabdomeric Eyes of Platynereis. Current Biology 25:2265–2271.

Huerta-Cepas J, Szklarczyk D, Heller D, Hernández-Plaza A, Forslund S, Cook H, Mende D, Letunic I, Rattei T, Jensen L, et al. 2019. eggNOG 5.0: a hierarchical, functionally and phylogenetically annotated orthology resource based on 5090 organisms and 2502 viruses. Nucleic Acids Research 47:D309–D314.

Katoh K, Rozewicki J, Yamada KD. 2017. MAFFT online service: multiple sequence alignment, interactive sequence choice and visualization. Briefings in Bioinformatics 20:1160–1166.

Kikuchi T, Dayi M, Hunt VL, Ishiwata K, Toyoda A, Kounosu A, Sun S, Maeda Y, Kondo Y, de Noya BA, et al. 2021. Genome of the fatal tapeworm Sparganum proliferum uncovers mechanisms for cryptic life cycle and aberrant larval proliferation. Communications Biology 4:649.

Kocot KM, Struck TH, Merkel J, Waits DS, Todt C, Brannock PM, Weese DA, Cannon JT, Moroz LL, Lieb B, et al. 2016. Phylogenomics of Lophotrochozoa with Consideration of Systematic Error. Systematic Biology 66:256–282.

Kück P, Longo GC. 2014. FASconCAT-G: extensive functions for multiple sequence alignment preparations concerning phylogenetic studies. Frontiers in Zoology 11:81.

Kumar S, Nei M, Dudley J, Tamura K. 2008. MEGA: A biologist-centric software for evolutionary analysis of DNA and protein sequences. Briefings in Bioinformatics 9:299–306.

Lan Y, Sun J, Chen C, Sun Y, Zhou Y, Yang Y, Zhang W, Li R, Zhou K, Wong WC, et al. 2021. Hologenome analysis reveals dual symbiosis in the deep-sea hydrothermal vent snail Gigantopelta aegis. Nature Communications 12:1165.

Lemoine F, Domelevo Entfellner JB, Wilkinson E, Correia D, Dávila Felipe M, De Oliveira T, Gascuel O. 2018. Renewing Felsenstein’s phylogenetic bootstrap in the era of big data. Nature 556:452–6.

Luo Y-J, Kanda M, Koyanagi R, Hisata K, Akiyama T, Sakamoto H, Sakamoto T, Satoh N. 2018. Nemertean and phoronid genomes reveal lophotrochozoan evolution and the origin of bilaterian heads. Nature Ecology & Evolution 2:141–151.

Maddison W, Maddison D. 2017. Mesquite: a modular system for evolutionary analysis. Version 3.31. http://www.mesquiteproject.org.

Marlétaz F, Peijnenburg KTCA, Goto T, Satoh N, Rokhsar DS. 2019. A New Spiralian Phylogeny Places the Enigmatic Arrow Worms among Gnathiferans. Current Biology 29:312-318.e313.

Martín-Durán JM, Vellutini BC, Marlétaz F, Cetrangolo V, Cvetesic N, Thiel D, Henriet S, Grau-Bové X, Carrillo-Baltodano AM, Gu W, et al. 2021. Conservative route to genome compaction in a miniature annelid. Nature Ecology & Evolution 5:231–242.

Minh BQ, Schmidt HA, Chernomor O, Schrempf D, Woodhams MD, von Haeseler A, Lanfear R. 2020. IQ-TREE 2: New models and efficient methods for phylogenetic inference in the genomic era. Molecular Biology and Evolution 37:1530–1534.

Morel B, Kozlov AM, Stamatakis A, Szöllősi GJ. 2020. GeneRax: A Tool for Species-Tree-Aware Maximum Likelihood-Based Gene Family Tree Inference under Gene Duplication, Transfer, and Loss. Molecular Biology and Evolution 37:2763–2774.

Passamaneck YJ, Furchheim N, Hejnol A, Martindale MQ, Lüter C. 2011. Ciliary photoreceptors in the cerebral eyes of a protostome larva. EvoDevo 2:6.

Picciani N, Kerlin JR, Sierra N, Swafford AJM, Ramirez MD, Roberts NG, Cannon JT, Daly M, Oakley TH. 2018. Prolific Origination of Eyes in Cnidaria with Co-option of Non-visual Opsins. Current Biology 28:2413-2419.e4.

Ramirez MD, Pairett AN, Pankey MS, Serb JM, Speiser DI, Swafford AJ, Oakley TH. 2016. The Last Common Ancestor of Most Bilaterian Animals Possessed at Least Nine Opsins. Genome Biology and Evolution 8:3640–3652.

Rangwala SH, Kuznetsov A, Ananiev V, Asztalos A, Borodin E, Evgeniev V, Joukov V, Lotov V, Pannu R, Rudnev D. 2021. Accessing NCBI data using the NCBI sequence viewer and genome data viewer (GDV). Genome Research 31:159–169.

Rawlinson KA, Lapraz F, Ballister ER, Terasaki M, Rodgers J, McDowell RJ, Girstmair J, Criswell KE, Boldogkoi M, Simpson F, et al. 2019. Extraocular, rod-like photoreceptors in a flatworm express xenopsin photopigment. eLife 8:e45465.

Seppey M, Manni M, Zdobnov EM. 2019. BUSCO: assessing genome assembly and annotation completeness with single-copy orthologs. Bioinformatics 31:3210–3212.

Serb JM, Eernisse DJ. 2008. Charting Evolution’s Trajectory: Using Molluscan Eye Diversity to Understand Parallel and Convergent Evolution. Evolution: Education and Outreach 1:439–447.

Shichida Y, Imai H. 1998. Visual pigment: G-protein-coupled receptor for light signals. Cellular and Molecular Life Sciences 54:1299–1315.

Shichida Y, Matsuyama T. 2009. Evolution of opsins and phototransduction. Philosophical Transactions of the Royal Society B 364:2881–2895.

Stanke M, Steinkamp R, Waack S, Morgenstern B. 2004. AUGUSTUS: a web server for gene finding in eukaryotes. Nucleic Acids Research 32:W309–W312.

Terakita A. 2005. The opsins. Genome Biology 6:213.

TheUniProtConsortium. 2020. UniProt: the universal protein knowledgebase in 2021. Nucleic Acids Research 49:D480–D489.

Vöcking O, Kourtesis I, Tumu SC, Hausen H. 2017. Co-expression of xenopsin and rhabdomeric opsin in photoreceptors bearing microvilli and cilia. eLife 6:e23435.

Westheide W, Rieger R. 2011. Zoologia sistematica. Filogenesi e diversità degli animali. Zanichelli.

Wollesen T, Rodríguez Monje SV, Oel AP, Arendt D. 2020. Characterization of eyes, photoreceptors and opsins in developmental stages of the chaetognath Spadella cephaloptera. bioRxiv:871111.

Yoshida MA, Ogura A, Ikeo K, Shigeno S, Moritaki T, Winters GC, Kohn AB, Moroz LL. 2015. Molecular Evidence for Convergence and Parallelism in Evolution of Complex Brains of Cephalopod Molluscs: Insights from Visual Systems. Integrative and Comparative Biology 55:1070–1083.

Zhang Y, Mao F, Mu H, Huang M, Bao Y, Wang L, Wong N-K, Xiao S, Dai H, Xiang Z, et al. 2021. The genome of Nautilus pompilius illuminates eye evolution and biomineralization. Nature Ecology & Evolution 5:927–938.

